# Encephalitic alphaviruses exploit caveolae-mediated transcytosis at the blood-brain barrier for CNS entry

**DOI:** 10.1101/805945

**Authors:** Hamid Salimi, Matthew D. Cain, Xiaoping Jiang, Robyn A. Roth, Wandy Beatty, Chengqun Sun, William B. Klimstra, Jianghui Hou, Robyn S. Klein

**Author notes:** Correspondence should be addressed to R.S.K. Robyn S. Klein, M.D., Ph.D., Washington University School of Medicine, Departments of Internal Medicine, Neurobiology, Pathology & Immunology, Campus Box 8051, 660 S. Euclid Ave, St. Louis, MO 63110.

## Abstract

Venezuelan and Western equine encephalitis viruses (VEEV and WEEV) invade the CNS early during infection, via neuronal and hematogenous routes (1, 2). While viral replication mediates host-shut off, including expression of type I interferons (IFN) (3, 4), few studies have addressed how alphaviruses gain access to the CNS during established infection or the mechanisms of viral crossing at the blood-brain barrier (BBB). Here, we show that hematogenous dissemination of VEEV and WEEV into the CNS occurs via caveolin (Cav)-1-mediated transcytosis (Cav-MT) across an intact BBB, which is impeded by IFN and inhibitors of RhoA GTPase. Use of reporter and non-replicative strains also demonstrates that IFN signaling mediates viral restriction within cells comprising the neurovascular unit (NVU), differentially rendering brain endothelial cells, pericytes and astrocytes permissive to viral replication. Transmission and immunoelectron microscopy revealed early events in virus internalization and Cav-1-association within brain endothelial cells. Cav-1-deficient mice exhibit diminished CNS VEEV and WEEV titers during early infection, whereas viral burdens in peripheral tissues remained unchanged. Our findings show that alphaviruses exploit Cav-MT to enter the CNS, and that IFN differentially restricts this process at the BBB.

**Importance:** VEEV, WEEV and EEEV are emerging infectious diseases in the Americas, and they have caused several major outbreaks in the human and horse population during the past few decades. Shortly after infection, these viruses can infect the CNS, resulting in severe long-term neurological deficits or death. Neuroinvasion has been associated with virus entry into the CNS directly from the blood-stream, however the underlying molecular mechanisms have remained largely unknown. Here we demonstrate that following peripheral infection alphavirus augments vesicular formation/trafficking at the BBB and utilizes Cav-MT to cross an intact BBB, a process regulated by activators of Rho GTPAses within brain endothelium. *In vivo* examination of early viral entry in Cav-1-deficient mice revealed significantly lower viral burdens than in similarly infected wild-type animals. These studies identify a potentially targetable pathway to limit neuroinvasion by alphaviruses.

## Introduction

The central nervous system (CNS) is protected from pathogens by the blood brain barrier (BBB), an intercellular association of transmembrane junctional proteins between brain microvascular endothelial cells (BMECs), with associated pericytes, astrocytes and neurons that together comprise the neurovascular unit (NVU) (5). Neurotropic pathogens have evolved mechanisms to bypass or cross this barrier, including anterograde or retrograde transport along axons, destabilization of BBB junctional proteins, or passage through BMECs (6), the latter of which may involve intracellular transport within leukocytes via binding to intercellular adhesion molecule (ICAM)-1 (7). An additional mechanism may involve caveolae, flask-shaped plasma membrane invaginations within BMECs that are important for cell metabolism, signal transduction, and the transcytosis of large proteins (8). BBB formation of caveolae, which contain the major structural protein caveolin-1 (Cav-1), is limited by the major facilitator superfamily domain-containing protein 2a (Mfsd2a), which is exclusively expressed on BMECs and induced by pericytes (9). Stabilization of junctional proteins and caveolae within BMECs is additionally regulated by the small Rho GTPases, including the Ras homolog gene family, member A (RhoA), and Ras-related C3 botulinum toxin substrate (Rac)-1 (10). While many viruses have evolved to interact with Rho GTPases to increase their entry and replication within target cells (11, 12), some neurotropic viruses, including retroviruses and flaviviruses, may compromise BBB permeability via GTPase-mediated alterations of junctional proteins (13–16). For these viruses, neuroinvasion coincides with BBB instability. In contrast, encephalitic alphaviruses, including Venezuelan, Western and Eastern equine encephalitis viruses (VEEV, WEEV and EEEV, respectively), can enter the CNS directly from the bloodstream via unknown mechanisms (17, 18).

VEEV, WEEV, EEEV naturally cycle between mosquitoes and birds (EEEV and WEEV), mosquitoes and rodents (VEEV enzootic cycle), or mosquitoes and horses (VEEV epizootic cycle), and are all widely distributed in North, Central, and South America (19). Human infection can progress rapidly to encephalitis with fatality rates of 1-75%, depending on the strain. Of the three, VEEV is considered the most important zoonotic pathogen with several reported outbreaks in South and Central Americas, the latter of which have spread to North America. Although the number of human cases reported is small, the possibility for disease emergence is high due to expansion and spread of mosquito vectors (20). Despite the epidemic potential of VEEV and the high morbidity and/or case fatality rates of EEEV and WEEV, there are no approved vaccines or therapeutics for humans. Insight into the cell-intrinsic and -extrinsic processes by which the host limits alphavirus infections and minimizes virus- and immune-induced injury is essential for developing strategies to contain virus dissemination and disease. While early studies suggested that VEEV enters the CNS via anterograde transport along peripheral nerves after cutaneous inoculation (1), recent findings, however, emphasize that VEEV may cross the BBB via unknown mechanisms (2, 21).

In this study, we show that peripherally inoculated virulent strains of VEEV and WEEV enter the CNS from the blood-stream as free virions through an intact BBB. While VEEV and WEEV interact, enter and traverse brain endothelium *in vivo*, virus replication within brain microvascular endothelial cells (BMECs) and pericytes is inhibited by type I IFN (IFN) signaling. Consistent with this, reporter and non-replicative strains of VEEV and WEEV were observed to first replicate within astrocytes and neurons, respectively, suggesting that alphaviruses cross the BBB without replication in BMECs or pericytes. Using an *in vitro* transcytosis assay, alphaviruses were found to utilize caveolin-mediated transcytosis (Cav-MT), which is directly regulated by small Rho GTPases, and, notably, IFN. Transmission and immunoelectron microscopy revealed *in vivo* virus encounter and entry at the BBB, with detection of VEEV within caveolin-1-expressing BMECs. Importantly, deficiency in *Cav-1* significantly delayed alphavirus neuroinvasion during early infection, highlighting the important role of caveolae in CNS entry. Together, these data suggest that IFNs regulate the CNS entry of encephalitic alphaviruses via direct and indirect mechanisms.

## Results

### Alphavirus neuroinvasion occurs prior to BBB disruption

To address mechanisms of alphavirus neuroinvasion in susceptible hosts, we infected mice with enzootic VEEV ZPC738 (VEEV) and epizootic WEEV McMillan (WEEV) strains of alphaviruses (22, 23). Wild-type mice infected with VEEV and WEEV via footpad (f.p.) inoculation exhibited detectable virus replication simultaneously in both fore- and hindbrain regions at 1- and 3-days post-infection (dpi), respectively (Fig. 1A and B). Similar results were observed in the brainstem and spinal cord (Fig. S1A-D). Consistent with prior studies, the olfactory bulb exhibited higher VEEV titers at early time-points (1). Viral titers plateaued at 10^7^ to 10^8^ pfu/g of tissue by 3-4 dpi (VEEV) and 4-5 dpi (WEEV). Significant alterations in BBB permeability occurred at plateau viral loads, with VEEV infection leading to higher permeability in all brain regions as compared to WEEV (Fig. 1C and D). Peak BBB permeability in the brainstem and spinal cord also occurred at time-points coinciding with peak viral loads (Fig. S1C and D). We also examined direct effects of virus on BBB integrity using an *in vitro* BBB model, in which primary murine BMECs are cultured on transwell inserts over primary murine astrocytes in the bottom chamber (Fig. 1E). In this model, barrier integrity is assessed by measuring transendothelial electrical resistance (TEER) between the transwell chambers. While control cultures treated with TNF-α (100 ng/ml) displayed decreased TEER at all time-points, addition of virus to either BMECs or astrocytes had no effect on TEER (Fig. 1F and G). Together, our observations from *in vivo* and *in vitro* experiments suggest that alphavirus neuroinvasion occurs in the presence of an intact BBB.

**FIG 1.**
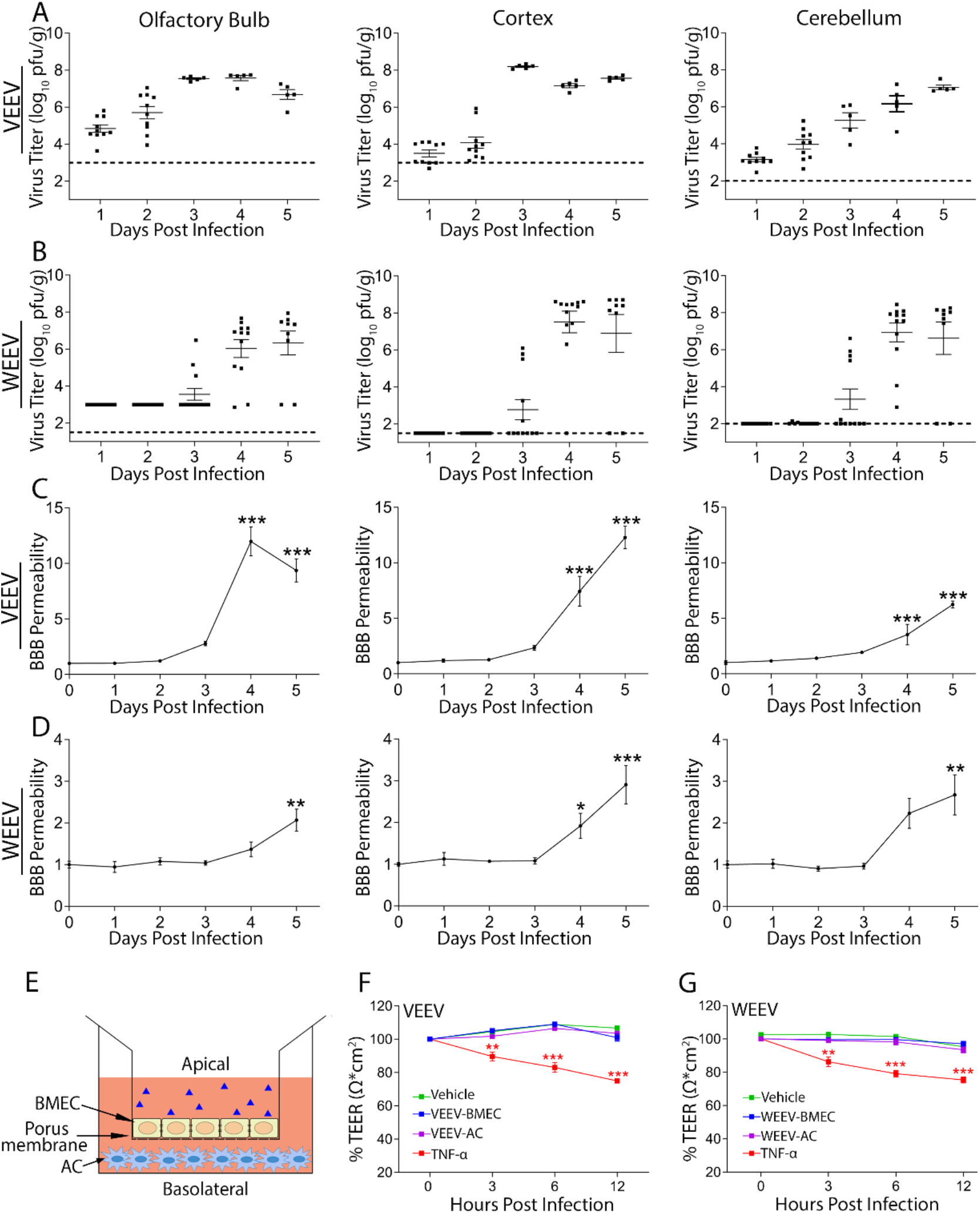
Alphavirus infection of the CNS occurs prior to BBB disruption. (A and B) Viral burdens in brain tissues of C57BL/6 mice following f.p. infection with either VEEV (10 pfu) or WEEV (1000 pfu), determined by plaque assay. Dashed lines indicate the limit of detection of the assay. Viral burdens were measured at 1 dpi (VEEV) and 3 dpi (WEE). (C and D) BBB permeability was determined at indicated dpi by measuring sodium fluorescein in CNS tissues following i.p. administration. Data presented as mean viral titer or fluorescence ± SEM for N=5-10 mice/group. *, *P* < 0.05, **; *P* < 0.01; ***, *P* < 0.001, via 1-way ANOVA. (E) A schematic figure of an *in vitro* BBB model. (F and G) Neither exposure of BMECs nor astrocytes (ACs) to VEEV and WEEV viruses had any effects on barrier integrity. Addition of TNF-α (100 ng/ml) significantly reduced TEER over time. Experiment was performed twice, each with 6 replicates. Error bars indicate mean ± SEM. Statistically significant differences were determined via 2-way ANOVA followed by Dunnett’s multiple comparison test.

### Hematogenous route of neuroinvasion is not exclusive to the circumventricular organs (CVOs)

Viral neuroinvasion across an intact BBB could occur via virus replication within cellular constituents of the NVU. To assess viral permissivity of NVU cells, we examined *in vitro* alphavirus infection of isolated murine cells and performed multistep growth curve analyses after infection with either VEEV (Fig. S2A-C) or WEEV (Fig. S2E-G). While all cell types were permissive to both viruses, slopes of the curves differed; thus, virus replication plateaued as early as 6-12 hours post-infection (hpi) in astrocytes, while pericytes and BMECs required 12-24 hpi and 24-48 hpi, respectively, to reach plateau levels. To validate these findings in murine BMECs, the experiment was performed using the human brain endothelial cell line, hCMEC/D3, which displayed similar results (Fig S1H and L). These data suggest that differential restriction of alphavirus replication may occur at the NVU. To examine this *in vivo* during early viral neuroinvasion, we infected mice with 10^6^ pfu of either VEEV-eGFP or WEEV-eGFP via intravenous (i.v.) injection, followed by immunohistochemical (IHC) detection of GFP within brain tissues. Given the two-day delay in detection of WEEV compared with VEEV within the CNS (Fig. 1) CNS tissues of VEEV-infected mice were examined at 16 hpi, while WEEV-infected mice were examined at 48 hpi. Analyses of brains revealed multiple foci of viral replication throughout the brain for both viruses, suggesting hematogenous routes of CNS entry. While some of the entry sites overlapped with areas consistent with CVOs (Fig. 2A and B; arrows) (2), others were located within cortical and cerebellar areas that are distant from these structures (Fig. 2A and B; arrow heads). In VEEV-infected mice, GFP expression was detected in both NeuN^+^ neurons and S100-β^+^ astrocytes (Fig. 2C). In contrast, WEEV-infected animals exhibited GFP expression exclusively within neurons (Fig. 2D). Notably, neither reporter strain led to early viral replication within BMECs or pericytes (Fig. 2C and D), which is consistent with their diminished permissivity *in vitro* compared with astrocytes. Together, these data suggest that cells of the NVU are differentially susceptible to alphavirus infection *in vivo* and that, upon viremia, alphaviruses invade the CNS from the bloodstream through multiple entry sites that are not exclusive to CVOs.

**FIG 2.**
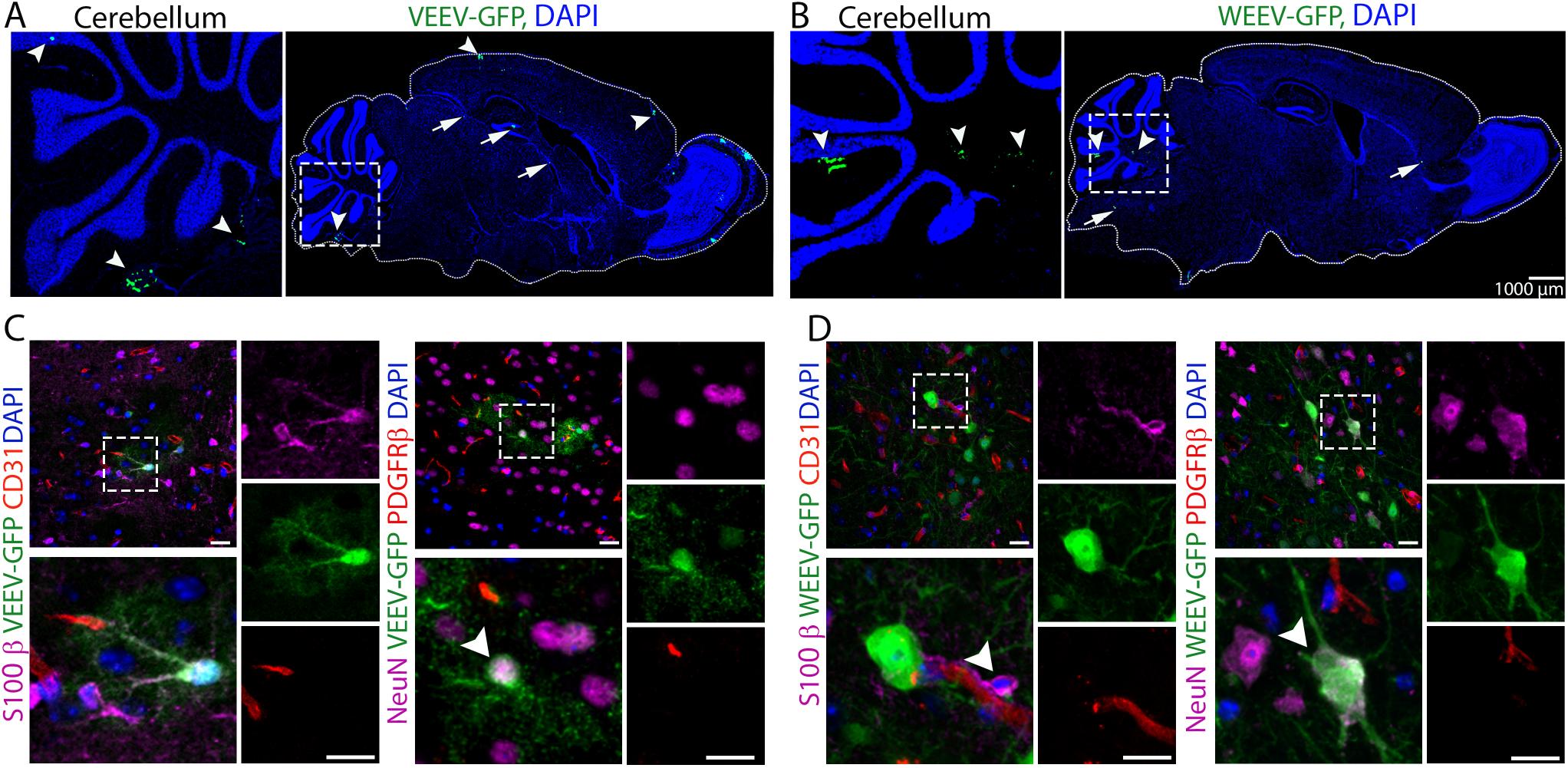
Hematogenous route of alphavirus neuroinvasion is not exclusive to CVOs. (A and B) Low magnification of a sagittal section of VEEV-eGFP- (A) and WEEV-eGFP (B) infected mouse brain showing multiple entry sites for CNS entry. Brain tissues from VEEV-eGFP (C) and WEEV-eGFP (D) infected mice were stained for makers of astrocytes (S100-β, Magenta), BMECs (CD31, Red), pericytes (PDGFR-β, Red) and neurons (NeuN, Magenta). Insets are enlarged in lower left of each panel. Single channels relate to the enlarged insets. Nuclei were counterstained with DAPI (blue). Magnification 40x, Scale bars: 20 µm.

### IFNAR signaling differentially restricts alphavirus infection at the BBB

Given our results demonstrating differential *in vivo* replication of alphaviruses within cells of the NVU, we hypothesized that robust post-entry restriction may be imposed by innate immune responses within BMECs and pericytes. To address this, WT and *Ifnar^-/-^*mice were infected with VEEV-eGFP (100 pfu) via f.p. inoculation, and brain tissues were examined at 1 dpi for GFP expression. Notably, *Ifnar^-/-^*mice succumb to VEEV-eGFP by ∼30 hr post infection (Fig. S3A), while WT animals have undetectable brain infection using IHC at this time-point (Fig S3D). Thus, to allow comparisons at similar viral burdens (Fig. S3C), we also evaluated brain tissues of WT mice at 3 dpi. VEEV infection was limited to S100-β^+^ astrocytes and NeuN^+^ neurons in WT animals (Fig. 3A and Fig. S4A), whereas similar infection in *Ifnar^-/-^* animals led to GFP expression within BMECs and pericytes both in the cortex (Fig. 3B) and cerebellum (Fig. S4B). Next, we assessed cellular tropism in the context of WEEV-eGFP (1000 pfu) infection. Although mortality was significantly increased in WEEV-eGFP-infected *Ifnar^-/-^*mice compared with similarly infected WT animals, *Ifnar^-/-^* mice survive WEEV infection up to 5-6 dpi (Fig. S3B), and viral infection was undetectable in brain tissues of either WT or *Ifnar^-/-^* mice at 1 dpi (Fig. S3E). Thus, brain tissues were examined at 3 dpi for both genotypes. Similar to our observations with VEEV, WT mice infected with WEEV exhibited no GFP expression in BMECs or pericytes, with infection detected only in neurons (Fig. 3C and Fig. S4C). However, *Ifnar^-/-^* mice additionally exhibited GFP expression within cortical (Fig. 3D) and cerebellar (Fig. S4D) astrocytes.

**FIG 3.**
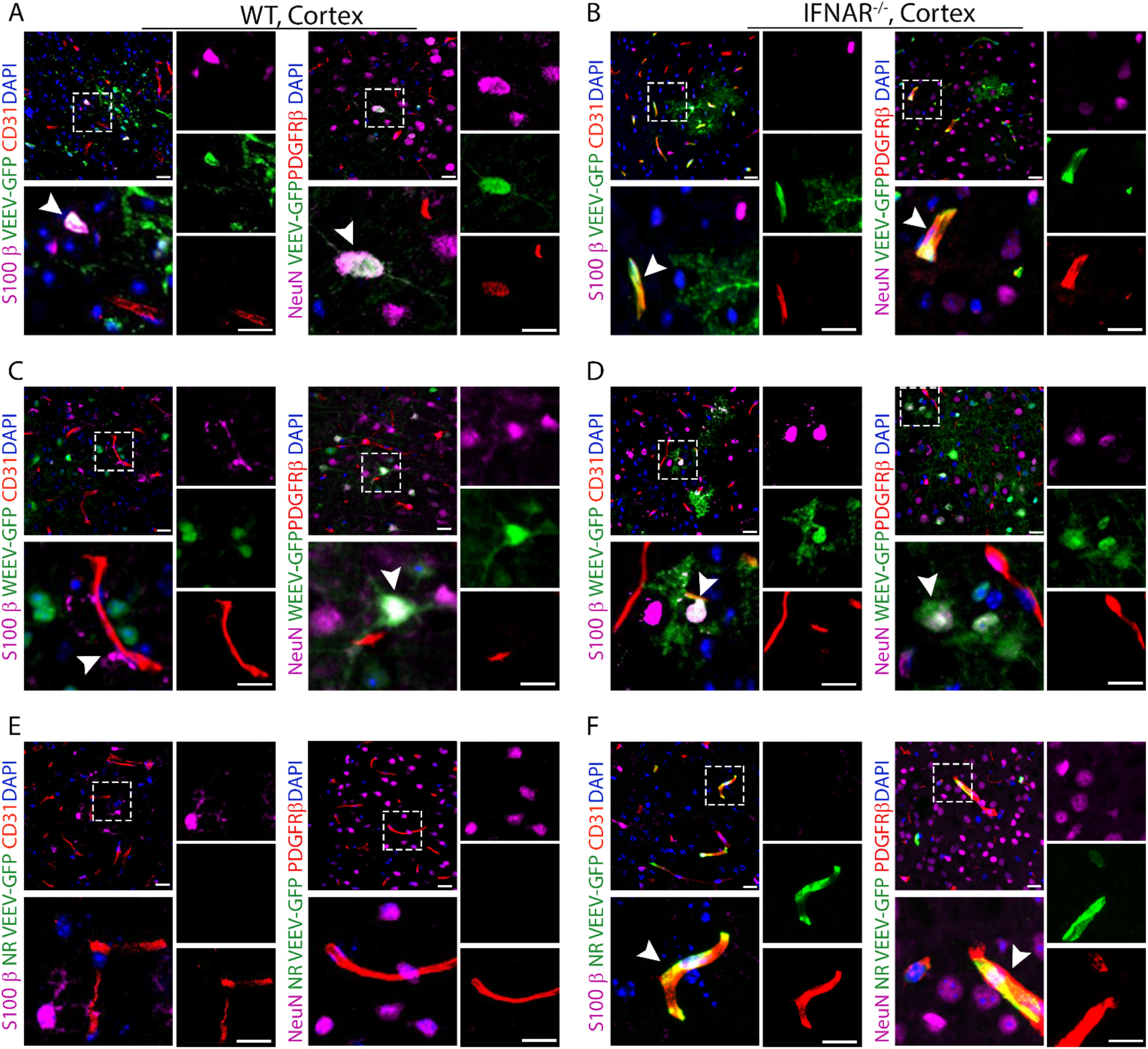
Type I IFNs differentially restrict alphavirus replication within the NVU. IHC staining of cortical brain regions of WT (A and C) and *Ifnar*^-/-^ (B and D) mice following f.p. infection with either VEEV-eGFP (A and B) or WEEV-eGFP (C and D). Cell markers: astrocytes (S100-β), BMECs (CD31), pericytes (PDGFR-β) and neurons (NeuN). Nuclei counterstained with DAPI (blue). (E and F) IHC staining of cortical brain regions of WT and *Ifnar*^-/-^ mice following i.v. infection with NR-VEEV-eGFP. Magnification 40x. Scale bars: 20 µm.

As the observed changes in cellular tropism could be due a higher titer of viremia in *Ifnar^-/-^* versus WT mice, *Ifnar^-/-^*mice were infected with 10^6^ pfu/ml of a non-replicative (NR) replicon derived from VEEV-eGFP (24); a physiologically relevant dose that is equivalent to the level of viremia observed in WT mice after f.p. inoculation (Fig. S4F). Following cellular entry, NR-VEEV-eGFP undergoes RNA replication and produces reporter genes products, but no progeny virus, due to lack of structural genes (24). Similar to our results with WT VEEV-eGFP, WT animals infected with NR-VEEV-eGFP had no infection of BEMCs or pericytes (Fig. 3E), whereas *Ifnar^-/-^*mice exhibited extensive infection of these cell types in multiple brain regions, including the cortex (Fig. 3F) and cerebellum (Fig. S4E). These findings support the notion that alphavirus infection is differentially restricted at the NVU by IFNAR signaling, which prevents CNS entry by blocking active virus replication within BMECs.

### Ultrastructural analysis reveals alphavirus interaction and entry at the BBB

To further characterize virus-cell interactions at the BBB *in vivo*, we performed transmission electron microscopy (TEM) of CNS tissues derived from VEEV-infected WT and *Ifnar^-/-^* mice. TEM analysis of *Ifnar^-/-^* mice, which develop high level viremia (Fig. S4F), revealed 70 nm spherical structures with electron dense cores, consistent with virions (25), attached to the luminal surface of cortical microvessels (Fig. 4A). Remarkably, immunogold labeling using antibodies against the VEEV E2 glycoproteins detected viral particles inside cortical BMECs of WT mice (Fig. 4B). Since WT BMECs do not support VEEV replication *in vivo* (Fig. 3A), detection of E2 glycoproteins likely represents an intact virion rather than intracellular protein expression. Based on these observations, we hypothesized that VEEV may cross the BBB as free particles. To examine this, mice were infected with NR-VEEV-eGFP via *i.v.* injection. At 1 dpi, we detected infection of astrocyte-like cells in multiple brain regions, including cortex and cerebellum (Fig. 4C and D), suggesting that VEEV can cross the BBB as free virions, likely via a transcytosis mechanism.

**FIG 4.**
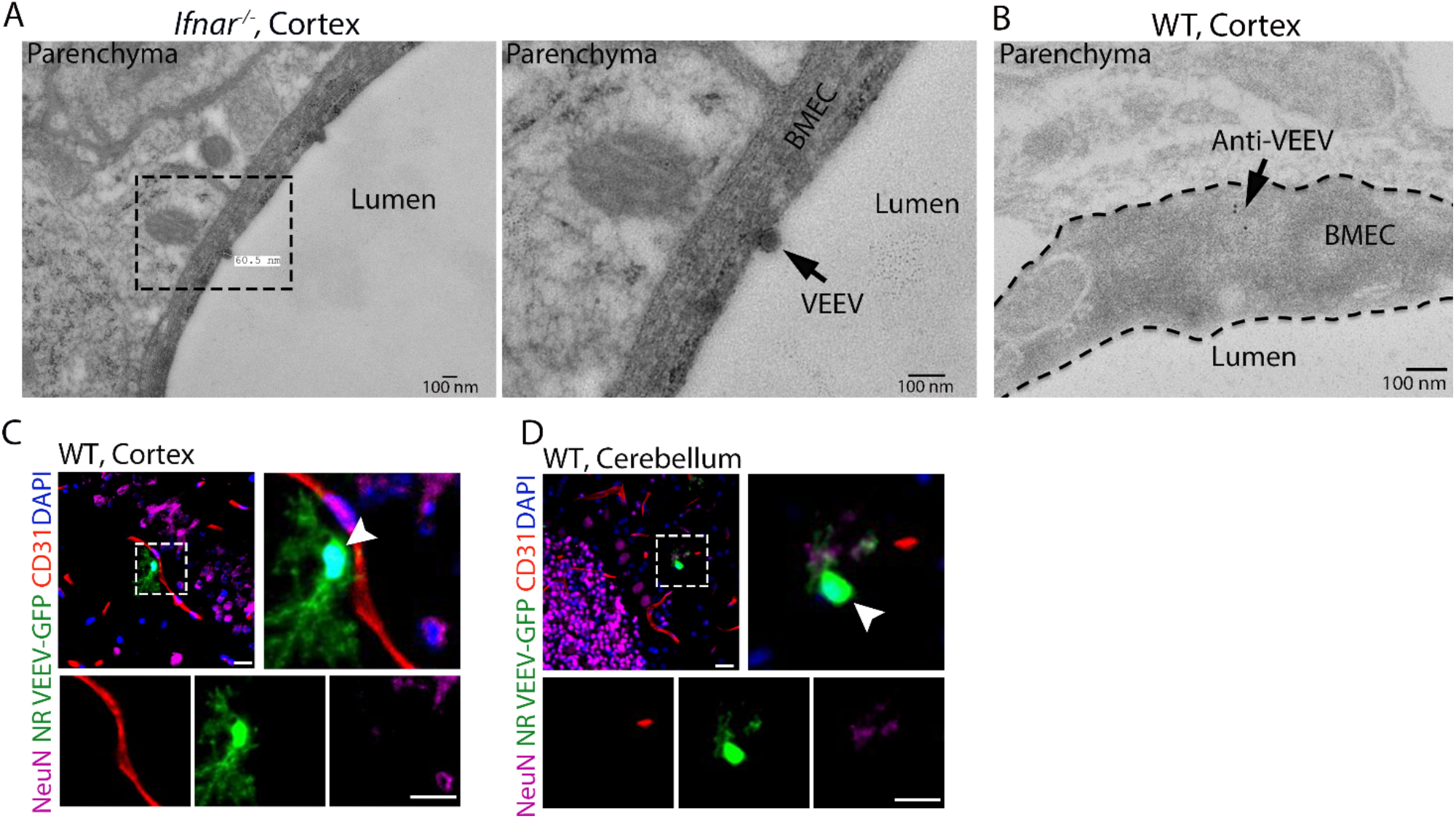
Detection of alphavirus interaction and entry at the BBB. (A) TEM analysis of brain tissues from VEEV infected mice at 1 dpi, identified virus like particles attached to luminal surface of microvessels. (B) Viral E2 glycoproteins were detected within cortical BMECs of VEEV-eGFP infected mice (3 dpi) by immune-EM using 8 nm immunogold particles. Scale bars=100 nm. (C and D) Immunostaining of brain tissues collected from NR VEEV-eGFP infected WT mice at 1 day following i.v. infection. Cell markers: BMECs (CD31), and neurons (NeuN). Nuclei counterstained with DAPI (blue). Magnification 40x. Scale bars: 20 µm.

### Alphavirus crosses brain endothelial cells via caveolae-mediated transcytosis

Based on our TEM data (Fig. 4A and B), we hypothesized that alphaviruses may exploit BMEC endocytic machinery for entry across the BBB. Contrary to peripheral tissues, the rate of transcytosis and numbers of endocytic vesicles are relatively low in BMECs (9). However, exposure of BMECs to VEEV or WEEV (MOI 10) resulted in enhanced caveolae formation at the cell surface (Fig. 5A-D). We next quantified the number of caveolae-like structures in BMECs of VEEV-infected mice. Similar to our *in vitro* data, we observed increased vesiculation in cytoplasm and at the luminal surface of BMECs in infected animals (Fig. 5E-G). In addition, virus infection induced membrane ruffling in BMECs of infected mice (Fig. 5H). These observations suggest a possible mechanism for viral transcytosis. To address this, we utilized a transcytosis assay in which virus is added to the top chambers of transwell inserts for 60 min, followed by removal of inserts and assessment of infectious virions in the bottom chamber (Fig. 6A). As the replication time of VEEV in BMECs is approximately 3-6 h (Fig. S5A), virus detected in the bottom chamber is unlikely due to replication, which was confirmed via assessment of GFP expression in bottom chamber astrocytes after transcytosis of NR-VEEV-eGFP virus (Fig. 6B). TEM evaluation of virus transcytosis across BMECs additionally identified virion-like structures within endosomes, characteristic of caveolae (26). Virion-containing endosomes were detected at various stages of transcytosis, i.e. early endosomes, multi-vesicular bodies and exocytic vesicles (Fig. 6C-E). Together, these data suggest that VEEV has the ability to cross endothelial cell barrier via a transcytosis mechanism.

**FIG 5.**
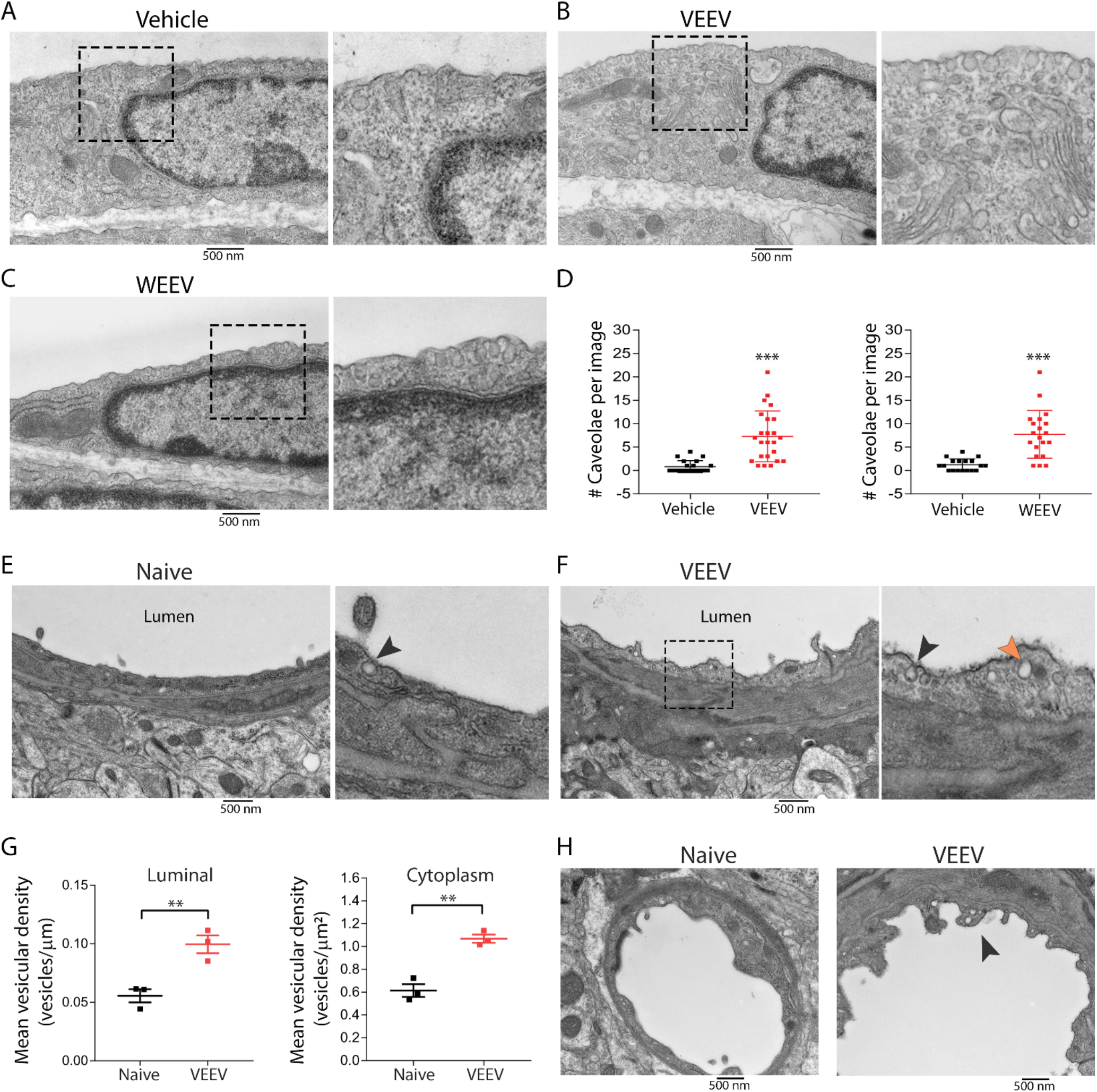
Alphavirus infection enhances caveolae formation in brain endothelial cells. (A-C) Exposure of BMECs to either VEEV or WEEV at a MOI of 10 enhanced caveolae formation at the cell surface compared to vehicle. (D) Quantitation of cell surface-associated caveolae in BMECs after virus exposure. Statistical differences were analyzed by 1-way ANOVA followed by Dunnett’s multiple comparison test. ***, *P* < 0.001. (E and F) TEM analysis of brain tissues collected from naïve and VEEV-infected mice at 3 dpi. (G) Quantitation of caveolae-like structures at the cell surface and inside cytoplasm of brain endothelial cells of naïve vs infected animals. Three tissue blocks were harvested from cortical brain region of each mouse (N=3 in each group), and 30 images were obtained per block. After manual counting, the mean vesicular density per length or volume of cytoplasm was calculated using ImageJ (NIH). Statistical differences were analyzed using t-test. (H) TEM analysis demonstrating membrane ruffling in BMECs of infected mice relative to control animals.

**FIG 6.**
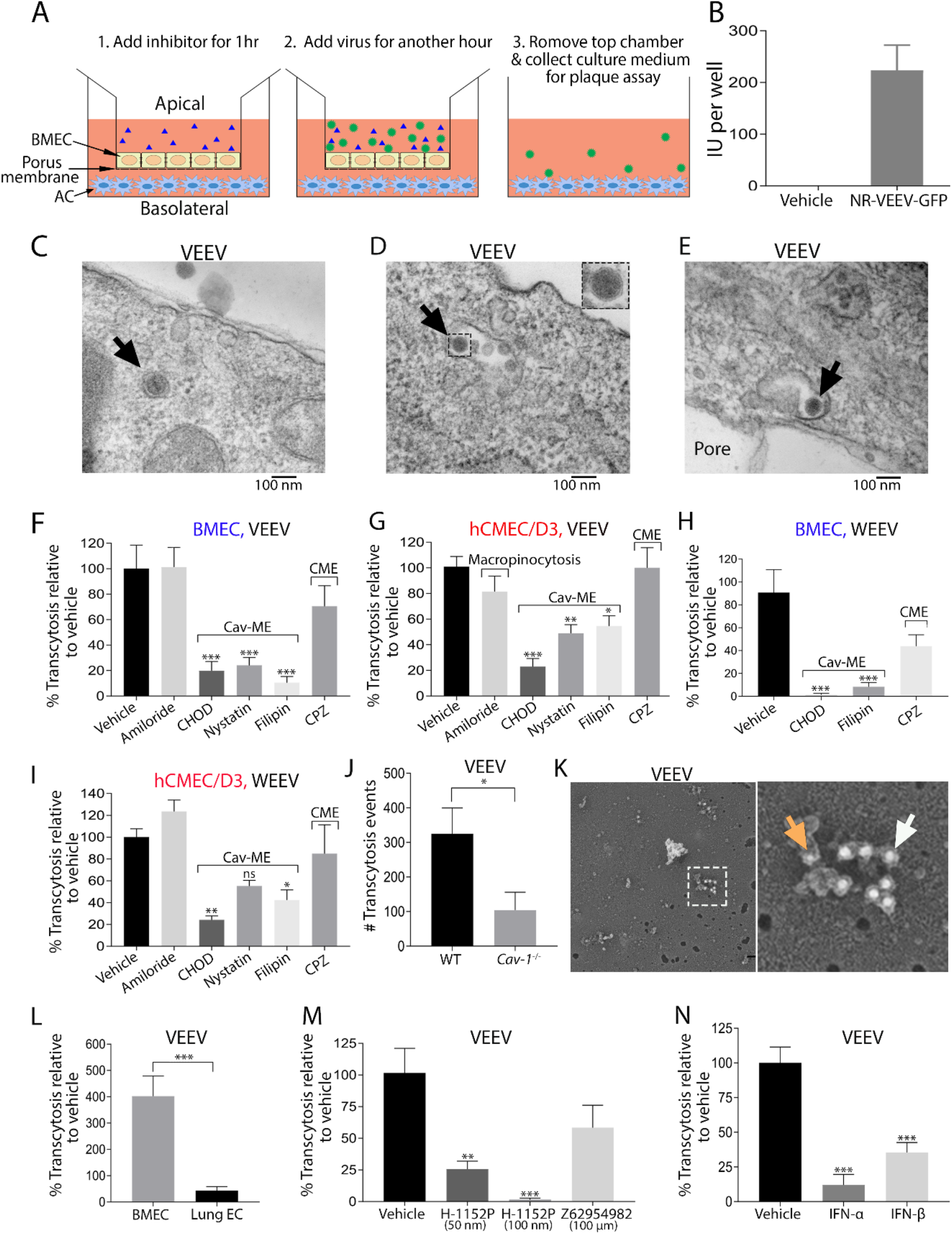
Alphavirus crosses BMECs via caveolae-mediated transcytosis. (A) A schematic figure demonstrating different steps of a transcytosis assay. (B) Addition of a non-replicative strain of VEEV-eGFP to hCMEC/D3 on the top chamber resulted in GFP signals in human astrocytes in bottom chamber. (C-E) TEM analysis identified virus like particles (depicted by arrows) within multi-vesicular bodies (D) and exotic endosomes (E) in BMECs after exposure to VEEV for 30 min. (F-I) VEEV and WEEV traverse across monolayers of BMEC (F and H) and hCMEC/D3 (G and I) via caveolae-mediated transcytosis. Data are presented as mean % of transcytosis relative to untreated cells. Experiments were repeated 2-3 times each with 4-6 technical replicates. Error bars indicate standard error of the mean (SEM). Concentration of inhibitors: Amiloride hydrochloride hydrate (50 µm), Cholesterol oxidase (CHOD, 2 U/ml), filipin (1 µg/ml), nystatin (12 µg/ml), chlorpromazine (CPZ, 10 µg/ml). (J) Quantitation of VEEV transcytosis across WT and *Cav-1*^-/-^ BMECs. (K) Immunogold labeling of virally infected BMECs revealed VEEV (8 nm immunogold particles; orange arrow) in colocalization with caveoline-1 (16 nm immunogold particles; white arrow). (L) VEEV transcytosis was compared between brain and lung ECs. (M and N) VEEV transcytosis across BMECs in the absence and presence of Rho kinase GTPase (H-1152P) and Rac1 (Z62954982), as well as type I IFNs (100 pg/ml). Results from transcytosis assays were analyzed by 1-way ANOVA (F-I, M and N), and t-test (J and L). *, *P* < 0.05, **; *P* < 0.01; ***, *P* < 0.001.

Transcytosis may utilize various pathways, including macropinocytosis, and clathrin- or caveolae-mediated endocytosis (CME & Cav-ME, respectively) (10, 27), which may be interrogated via use of inhibitors including amiloride hydrochloride (macropinocytosis); filipin, nystatin and cholesterol oxidase (Cav-ME), and chlorpromazine (CME). Compared with untreated BMECs or hCMEC/D3, the percentage of transcytosis for both VEEV and WEEV was significantly reduced in the presence of inhibitors of Cav-ME, but not CME or macropinocytosis (Fig. 6F-I). Notably, none of the inhibitors affected viral infectivity or barrier integrity (Fig. S5B-E). In accordance with the key role of caveolin-1 in caveolae formation, *Cav-1*^-/-^ BMECs displayed diminished virus transcytosis as compared to WT cells (Fig. 6J). These results were further verified using freeze fracture electron microscopy, wherein we identified VEEV particles (8 nm gold particle; orange arrow) co-localized with Cav-1^+^ (16 nm gold particles, white arrow) (Fig. 6K). In addition, lung endothelial cells exhibited significantly reduced virus transcytosis compared to BMECs (Fig. 6L), suggesting specificity of alphavirus transcytosis for BMECs.

Caveolae formation is enhanced by the small Rho GTPase RhoA, which is inhibited by Rac1 activity (28, 29). Consistent with this, H-1152 (Rho-kinase inhibitor) significantly reduced VEEV transcytosis across BMECs, while Z62954982 (Rac-1 inhibitor) had no significant effect on virus transmigration (Fig. 6M). However, treatment with IFN-α or IFN-β, which has been shown to promote BBB tight junction integrity via activation of Rac-1 (15), significantly reduced VEEV transcytosis, suggesting direct effects of IFNAR signaling on Cav-MT (Fig. 6N). Together, these results indicate that VEEV and WEEV exploit Cav-1-MT to cross an intact BBB and that this may be regulated by IFN.

### Caveolin-1 contributes to alphavirus neuroinvasion *in vivo*

To validate the role of caveolin-1 in early alphavirus neuroinvasion *in vivo*, we infected WT and *Cav-1^-/-^* mice with either VEEV (10 pfu) or WEEV (1000 pfu) via f.p. infection and examined CNS viral burdens at 1 and 3 dpi. Both VEEV and WEEV-infected *Cav-1*^-/-^ mice exhibited significantly reduced viral titers in the cortex and cerebellum at 1 dpi and 3 dpi, respectively, as compared to similarly infected WT animals, whereas viral burden in the serum and spleen were indistinguishable between the two genotypes (Fig. 7A and B). Importantly, no differences in CNS viral loads were detected in intracranial alphavirus-infected WT versus Cav-1^-/-^ mice (Fig. S6A and B), indicating no effect of Cav-1-deficiency on viral replication within the CNS. These results indicate that caveolin-1 critically contributes to alphavirus neuroinvasion after peripheral infection.

**FIG 7.**
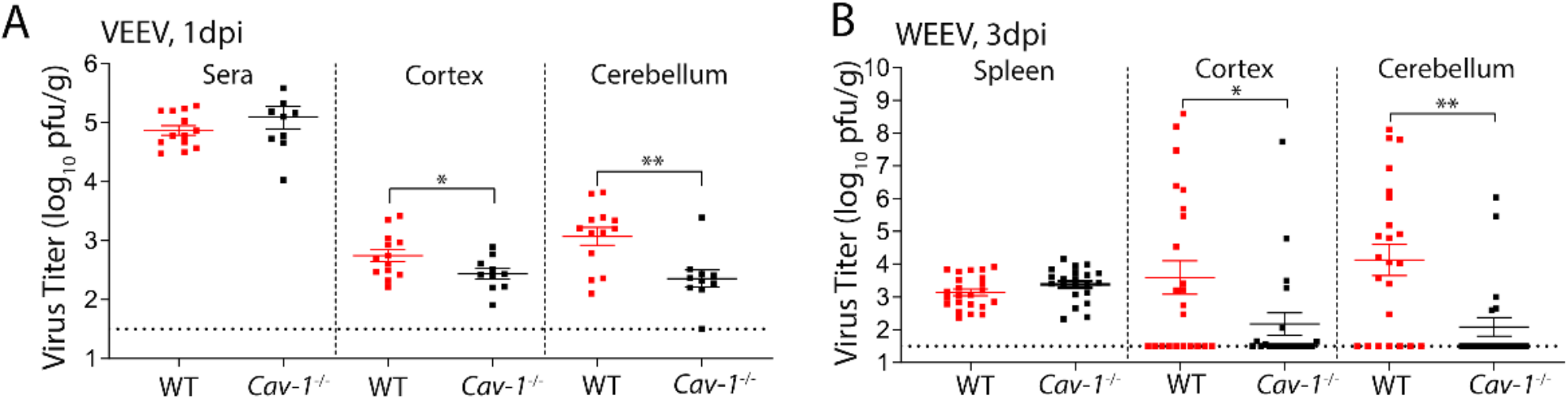
*Cav-1^-/-^* mice displayed reduced viral titers in the brain as compared to WT animals. (A and B) Viral burdens in peripheral and brain tissues of WT vs Cav-1 KO mice after f.p. infection with VEEV (10 pfu) and WEEV (1000 pfu), determined by plaque assay. Error bars indicate standard error of the mean (SEM). Shown is the combined data from 4 independent experiments each with 4-6 animals. Results were analyzed by unpaired t-test. *, P < 0.05; **, P < 0.01.

## Discussion

Using *in vitro* and *in vivo* approaches, we examined early events during alphavirus neuroinvasion. We found that VEEV and WEEV can enter the CNS via hematogenous dissemination across an intact BBB, without viral replication within BMECs or pericytes, leading to productive infection of CNS resident cells. Differential restriction of viral infection within cellular constituents of the NVU is mediated by IFNAR signaling, with replication in BMECs and pericytes (VEEV), and astrocytes (WEEV) only observed in *Ifnar^-/-^*mice. VEEV and WEEV entry across the BBB occurs via Cav-MT, which is impeded by Rho kinase inhibitor and IFN, the latter likely via activation of Rac1 (15, 16). Immuno-EM demonstrated alphavirus interaction, internalization and Cav-1 association within BMECs both *in vitro* and *in vivo*. Consistent with this, Cav-1-deficiency reduced alphavirus transcytosis *in vitro* and led to reduced titers of VEEV or WEEV in the CNS during early infection *in vivo*. Taken together, these data suggest that innate immune signaling regulates alphavirus neuroinvasion and replication at the NVU.

The CNS entry of alphaviruses following peripheral infection has been shown to occur via multiple routes. Early studies in mice indicated that VEEV may reach the brain via retrograde transport along peripheral nerves (1), whereas, more recently VEEV and other encephalitic alphaviruses have been shown to enter the CNS directly from the bloodstream (2, 17, 30). In our study, peripheral infection of mice with virulent strains of VEEV and WEEV resulted in detectable viral loads at time-points that precede BBB disruption (Fig. 1), and neither VEEV nor WEEV induced alterations in TEER across an *in vitro* BBB. These results indicate that alphaviruses cross the BBB shortly after infection without affecting barrier integrity. Hematogenous routes of neuroinvasion may include viral passage through the CVOs, which lack specializations that comprise the BBB (2). Our study demonstrated that shortly after *i.v.* infection (16hrs), foci of alphavirus infection are observed simultaneously in cortical and cerebellar locations distant from the CVOs, suggesting that CNS entry is not exclusive to these structures. Notably, BBB breakdown occurs at later time-points, when viral loads have peaked, likely due to induction of inflammatory responses and infiltration of immune cells into the CNS (21, 31, 32).

Pathogens, including viruses, may cross the BBB via direct infection of brain endothelium (33, 34). Use of reporter and non-replicative strains of VEEV and WEEV during *in vivo* infection, however, indicated lack of infection within BMECs (Fig. 3), suggesting that active virus replication in brain endothelium does not contribute to CNS entry. VEEV has a broad tissue tropism and can infect many cell types, including macrophages and dendritic cells (24). Infected leukocytes may act as Trojan horses and introduce viral particles into the CNS upon infiltration. However, leukocyte interaction and extravasation at the BBB require elevated expression of cell adhesion molecules (CAMs) on brain endothelium, which does not occur until 3dpi (21). Since we detected significant viral loads throughout the CNS by 1 dpi (Fig. 1), it is unlikely that infected leukocytes are the initial source of virus in the CNS. Importantly, our data using NR-VEEV-eGFP did not detect viral infection within CNS infiltrating leukocytes, suggesting that virus can enter CNS as free virion and independent of leukocyte trafficking. Nonetheless, additional studies are required to precisely define the role of Trojan horse in alphavirus neuroinvasion.

Our TEM analysis demonstrated virus like particles attached to the lumen of brain endothelium. Given that viremia arises as early as 8 hr post VEEV infection, BMECs are likely among the first cell types exposed to virions. Such exposure may trigger innate immune responses (e.g. type I IFNs), which can influence intercellular communication at the NVU (35). Indeed, in the context of IFNAR deficiency, VEEV and WEEV replicate within BMECs and astrocytes, respectively. The lack of viral replication within these cellular constituents of the NVU in WT animals is likely due to paracrine effects of IFN that induce antiviral proteins, as has been recently reported for arthritogenic alphaviruses (36, 37). VEEV attachment to and internalization within BMECs was further confirmed via immuno-TEM, in which viral E2 glycoproteins were detected within these cells, supporting the notion that alphaviruses may cross the BBB via a transcytosis pathway.

Compared to peripheral tissues, the rate of transcytosis at the BBB is unusually low. However, this rate may be augmented by increased Src kinase activity, which is mediated by a group of pathologic and non-pathologic stimuli, including inflammatory mediators and immune cell interactions with brain endothelium (38). Viruses may also increase vesicular trafficking, as has been observed in peripherally-derived endothelial cells exposed to dengue virus (39). In our study we utilized both *in vitro* assays and *in vivo* viral infection models, and observed that both VEEV and WEEV induce formation of caveolae in BMECs. *In vivo*, VEEV infected mice exhibited increased vesiculation and membrane ruffling within brain endothelial cells (Fig. 5). *In vitro* studies have shown that alphaviruses may induce dramatic structural changes in the actin cytoskeleton, leading to the formation of filopodia-like extensions in infected cells (40). Notably, expression of the viral non-structural protein (nsP)-1 alone is sufficient to trigger formation of short extensions, which is dependent on its palmitoylation activity (41). Consistent with this, ablation of nsP1 palmitoylation sites abolishes the ability of Semliki Forest virus (SFV) to infect the brain in murine models (42). Elucidating molecular mechanisms underlying alphavirus induced vesiculation in BMECs may reveal therapeutic targets against viral neuroinvasion, and provide insights for BBB maintenance in the context of other neurological disorders induced by increased transcytosis in brain endothelium (43–45).

Transcytosis of macromolecules across brain endothelium predominantly occurs via mechanisms that utilize caveolae. During infectious diseases, interaction with caveolae allows pathogens to escape lysosomal degradation and cross endothelial cell barriers (46, 47). While different families of viruses have been shown to cross epithelial and endothelial cell barriers utilizing Cav-MT *in vitro* (48–50), *in vivo* findings to support these data have been difficult to obtain, especially within the brain where these events are rare. Our study utilized multiple approaches to demonstrate that neurotropic alphaviruses enter but do not replicate within brain endothelium, and that Cav-MT contributes to *in vivo* viral infection of the CNS. As peripherally infected *Cav-1*^-/-^ mice exhibit a delay in achievement of peak brain titers of VEEV and WEEV compared to WT animals, entry of virus in the absence of caveolae may rely on slower mechanisms of viral entry, such as retrograde transport along axons (1). Additionally, ablation of caveolin-1 results in upregulation of caveolin-independent pathways (51), which might explain how viral loads within CNS tissues derived from *Cav-1^-/-^*mice quickly attain the levels observed in WT animals.

The signaling cascade underlying transcytosis was further elaborated using *in vitro* assays, wherein deficiency in caveolin-1 or depletion of membrane cholesterol significantly reduced VEEV and WEEV transcytosis across BMEC monolayer. Notably, virus transcytosis across brain endothelial cells was blocked by Rho kinase inhibitor (Fig. 6). The Rho family of GTPases, including RhoA, Rac-1 and Cdc42 are known to play a role in pathogen uptake and dissemination within the host (52–55). These proteins alternate between an active GTP-bound, and an inactive GDP-bound form, which triggers rearrangements in the actin cytoskeleton and cellular uptake of pathogens (56). Studies in flaviviruse encephalitis have shown that IFNβ acts in synergy with the TAM receptor Mertk to activate of Rac-1, which enhances BBB tight junction integrity, thereby preventing paracellular entry of virus (15, 16). In the current study, we discovered that type I IFN also prevents alphavirus transcytosis across BMECs. Thus, although the VEEV nsP2 and capsid protein inhibits IFN signaling via host-shut off (4), the lack of viral replication within BMECs may allow IFN signaling to limit viral entry likely via modulation of Rho GTPases (15).

In summary, mechanisms of encephalitic alphavirus neuroinvasion from the blood are complex and may depend on individual viral tropism for olfactory sensory neurons and cellular constituents of the NVU, which provide avenues of entry that cross the BBB, respectively (24, 57). Neuroinvasion may also depend on innate immune mechanisms that exert virus and cell-specific effects on replication (4). Ablation of the olfactory route reduces but does not eliminate CNS entry of VEEV (1), suggesting each route of entry provides a relative contribution to alphavirus neuroinvasion. Further studies are needed to address how each process individually contributes to viral entry, and whether viral spread within the CNS relies on these pathways or on innate immune mechanisms that regulate intracellular viral trafficking and replication within permissive cells. Our data highlighted the critical role of Cav-MT in alphavirus neuroinvasion. Delineating molecular mechanisms involved in this process may reveal novel targets for potential therapeutic interventions against CNS infection.

## MATERIALS AND METHODS

### Cells

Baby hamster kidney (BHK-21) and African green monkey kidney (Vero) cells were maintained in Dulbecco’s Modified Eagle’s medium (DMEM) supplemented with 10% fetal bovine serum (FBS) and 100 µg/ml of penicillin and streptomycin. Primary murine brain microvascular endothelial cells (BMECs), pericytes and astrocytes were isolated from cortical brain of C57BL/6J mice, and maintained in culture as described previously (15, 58). The immortalized human cerebral microvascular endothelial cell line hCMEC/D3 (59) was purchased from Millipore Sigma and maintained in Endothelial Cell Growth Basal Medium-2 (EBM2, Lonza). Primary human astrocytes and murine lung endothelial cells were purchased from Sciencell Research Laboratories and Cell Biologics, respectively. Cells were maintained in the culture medium recommended by each company.

### Antibodies and reagents

Rabbit polyclonal anti-S100-β (ab41548), rabbit monoclonal anti-Calbindin (ab108404) and rabbit polyclonal anti-RFP (ab62341) antibodies (Abs) were purchased from Abcam Biotechnology. Purified rat anti-mouse CD31 (550274, clone MEC 13.3) and guinea pig anti-NeuN (ABN90P) polyclonal Abs were purchased from BD Biosciences and Millipore-Sigma, respectively. Goat anti-PDGFR-β (AF1042) Ab was purchased from R&D Systems. The anti-VEEV E2 monoclonal antibody was a kind gift from Dr. Michael Diamond (Washington University in St Louis). Amiloride hydrochloride hydrate, filipin, cholesterol oxidase, nystatin, chlorpromazine hydrochloride, Z62954982 and fluorescein sodium salt were purchased from Sigma-Aldrich. (S)-Glycyl-H-1152 (hydrochloride) was purchased from Cayman company. Mouse IFN-α and -β were obtained from PBL Assay Science.

### Virus propagation, purification, and titration

All viruses used in this study including the non-replicative “replicon” strain of VEEV (NR-VEEV-eGFP), replication competent VEEV, McMillan (WEEV), and GFP reporter viruses (60) were generously provided by Dr. William Klimstra (University of Pittsburgh, Pittsburgh, PA). The parental VEEV and WEEV cDNAs were gifts to Dr. Klimstra from Dr. Scott Weaver, University of Texas Medical Branch at Galveston and Dr. Kenneth Olson, Colorado State University, respectively. The McMillan cDNA was modified by placing the entire virus coding region in a pBR322-based plasmid under control of at T7 bacteriophage promoter. Stocks of VEEV and WEEV viruses were generated as described previously (61). Briefly, BHK-21 cells were infected at a multiplicity of infection (MOI) of 0.1. Culture medium was replaced with fresh medium 4 hr later. Virus-containing supernatants were collected on the next day (∼30 h post infection), cleared from cell debris by low-speed centrifugation and filtered through 0.22 μm filters. Virus particles were then concentrated by ultracentrifugation at 100,000 g for 2 h at 4°C through a cushion of 30% (wt/wt) sucrose in PBS. The virus pellet was re-suspended in PBS and stored in single use aliquots at −80°C. Titration of VEEV and WEEV were performed in BHK-21 and Vero cells, respectively.

### Multi-step growth curves

Primary murine BMECs, astrocytes and pericytes were seeded in 24-well plates for 3-4 days till they reached confluence. At this point cells were exposed to MOIs of 0.01, 0.1, and 1 of VEEV and WEEV viruses. Culture medium was removed one hour post infection and cells were washed 4 times with 1 ml/well of PBS. The last wash was stored at −80°C, and later it was used in plaque assay to determine residual unbound virus remaining in each well. Culture supernatant was harvested at specified time points and virus titer was determined using plaque assay in BHK-21 (VEEV) and Vero cells (WEEV).

### Mice studies

C57BL/6J wild-type and caveoline-1 knockout mice were purchased from Jackson Laboratory (Bar Harbor, ME). *Ifnar*^-/-^ mice were kindly provided by Dr. Michael Diamond (Washington University School of Medicine, St. Louis, MO). Animals were housed under pathogen-free conditions, and the experimental procedures were completed in accordance with the Washington University School of Medicine Animal Safety Committee. Male, 8- to 9-week-old mice were used in all *in vivo* experiments. Mice infections were performed by administrating virus either subcutaneous (SC), intravenous (IV) or intracranial (IC) injection, while the mice were under light ketamine anesthesia. For subcutaneous infection, mice were injected in the footpad with VEEV (10 pfu) and WEEV (1000 pfu) in 50 µl PSB. Intracranial infections were performed by inoculating 10 pfu (VEEV) or 100 pfu (WEEV) of virus in 10 µl of PBS into the right cerebral hemisphere via a guided 29-gauge needle. Intravenous infections were achieved by administrating 2*10^6^ pfu of virus in 100 µl of PBS via retro-orbital injection. Mock infected animals received the same volume of PBS in each infection method and were considered as controls where needed.

### Measurement of viral burden in tissues

To monitor the kinetic of virus spread *in vivo*, peripheral organs and CNS tissues were harvested from infected mice at specified time points. Prior to this, mice were perfused transcardially with 30 ml PBS. Tissues were homogenized in 500 µl PBS using MagNA Lyser (6000 rpm for 1 min) instrument, and virus titer in tissues homogenates was determined using plaque assay in either BHK-21 or Vero cells depending on virus. Alternatively, groups of mice were followed for survival assays.

### *In vivo* assessment of BBB permeability

At given days post infection, mice received 100 µl of 100 mg/ml fluorescein sodium salt (NaFL) in PBS via intraperitoneal injection. When the salt has reached equilibrium (45 min), blood samples were collected, and mice were perfused transcardially with 30 ml PBS prior to collection of CNS tissues. Serum samples and tissue homogenates were treated with 2% trichloroacetic acid overnight at 4°C to precipitate proteins. Supernatants were clarified from cellular debris by centrifugation (4000 rpm for 20 min at 4°C) and diluted in equal volumes of borate buffer, pH 11 (Sigma-Aldrich). Concentration of NaFl in supernatants was determined by measuring fluorescence emission at 538 nm using Synergy H1 microplate reader (BioTek Instruments, Inc.). Measurement values were normalized to tissue weight and to NaFL plasma concentration in each mouse.

### *In vitro* BBB model and transcytosis assay

An *in vitro* BBB model was generated as described elsewhere (61). Briefly, BMECs were seeded on the apical side of fibronectin-coated inserts (BD Falcon, 24-well, 3 µm pores). Concurrently, primary murine astrocytes were cultured in fibronectin-coated 24-well plates. Two days later, when astrocytes reached confluent, BMECs inserts were moved to astrocytes-containing plate. Astrocytes release growth factors that promote barrier formation between BMECs. Culture medium was changed every 3 days. On day 7, hydrocortisone (550 nM), CTP-cAMP (250 µM) and RO 20-1724 (17.5 µM) compounds were added to the cells in serum free medium to further promote expression of tight junctions and barrier formation between BMECs. On the following day, barrier integrity was assessed by measuring transendothelial electric resistance (TEER) before cells were used for transcytosis assay. In these assays, BMECs on the top chamber are exposed to different treatments for 1-2 h. Virus particles are then added at a MOI of 2 to BMECs on the top chamber for an additional hour. At this point, BMEC inserts are removed and culture medium is collected from the bottom chamber and is used in plaque assay to determine the number of virus particles that have crossed the BMECs monolayer in the presence and absence of different inhibitors. Exposure of BMECs to alphavirus is limited to one hour to prevent active virus replication and release of viral progenies from the basolateral side.

### Immunohistochemistry and confocal microscopy

Brian tissues were collected from infected mice at specified time points as indicated in each figure legend. Prior to tissue collection, mice were deeply anesthetized and perfused transcardially with 30 ml of PBS followed by 30 ml of 4% paraformaldehyde (PFA). Tissues were stored overnight in 4% PFA in PBS at 4□C, then transferred into two exchanges of 30% sucrose for 48 h, before they were embedded in O.C.T compound (Tissue-Tek). 10 µm cryostat brain sections were treated with proteinase K (5 µg/ml for 30 min at RT) for antigen retrieval. After 30 min incubation in blocking buffer, tissues sections were stained with primary antibodies to markers for astrocytes (S100 calcium-binding protein (S100)-β), pericytes (platelet-derived growth factor receptor beta/PDGFR-β), BMECs (CD31) and neurons (NeuN and calbindin). Sections were then incubated with appropriate secondary antibodies (Alexa Fluor 488, 555, and Dylight 650, all from Invitrogen) for 15 min in blocking buffer, followed by three washes in PBS. After antibody labelling, cells were counterstained with DAPI (D1306, Invitrogen). All images were obtained using confocal microscope (Carl Zeiss), and processed with ImageJ software (NIH).

### Transmission- and immunogold electron microscopy

For TEM, mice underwent cardiac perfusion with 5 ml of PBS, followed by 20 ml of 1.5% glutaraldehyde and 1% PFA in 0.12 M sodium pyrophosphate buffer, while for immuno-EM studies, perfusion was performed with 5 ml of PBS and 20 ml of 4% paraformaldehyde in 0.12 M sodium pyrophosphate buffer. Brain tissues were then collected and stored in fixative buffer overnight at 4°C and processed as previously described(61). For immuno-EM, ultrathin brain sections (70 nm) were stained with primary and gold-conjugated secondary antibodies on carbon-coated glass. Sections were viewed on a JEM-1400 transmission microscope (JEOL) at 80 KV with an AMT XR111 4k digital camera. For *in vitro* assays, monolayer of BMECs on Transwell inserts was infected with alphavirus at a MOI of 200. After 30 min, culture medium was replaced with fixative buffer (2% PFA and 2.5% glutaraldehyde in 100 mM sodium cacodylate buffer, pH 7.2) and incubated for 1 h at room temperature. Cells were then rinsed with sodium cacodylate buffer, and embedded in a thin layer of 2.5% agarose, followed by 1 h of fixation in 1% osmium tetroxide (Polysciences Inc.). After extensive wash with dH_2_0, cells were stained with 1% aqueous uranyl acetate (Ted Pella Inc., Redding, CA) for 1 h, dehydrated through a series of ethanol concentrations in distilled water, and embedded in Eponate 12 resin (Ted Pella Inc.). Ultrathin sections were generated at 95 nm using UCT ultramicrotome (Leica Microsystems Inc., Bannockburn, IL), and stained with uranyl acetate and lead citrate. Electron micrographs were obtained using a transmission electron microscope JEOL 1200 EX (JEOL USA Inc., Peabody, MA) equipped with an AMT 8-megapixel digital camera and AMT Image Capture Engine V602 software (Advanced Microscopy Techniques, Woburn, MA).

### Freeze fracture deep etching electron microscopy

Freeze fracture electron microscopy was achieved as described periviously(62). Briefly, cultured BMECs were rapidly frozen by abrupt application of the sample against a liquid helium-cooled copper block with a Cryopress freezing machine. Samples were then moved to a liquid nitrogen-cooled Balzers 400 vacuum evaporator, fractured, and etched at −104°C for 2.5 min. For immuno-freeze fracture EM, etched samples were first stained with primary and gold-conjugated secondary antibodies on carbon-coated glass. These samples were then rotary replicated with platinum (∼2 nm), deposited from a 20° angle above the horizontal plane, followed by an immediate, stabilization film of pure carbon (∼10 nm) deposited from an 85° angle. Replicas were floated onto bleach and transferred through multiple rinses of dH2O before placing on formvar-coated EM grids. Electron micrographs were obtained using a JEM1400 transmission microscope (JEOL) at 80 KV, equipped with an AMT XR111 4k digital camera.

### Statistical analysis

Statistical analysis was performed using GraphPad Prism 7 software. A probability value of p < 0.05 was considered statistically significant. Statistical values are indicated as *, P < 0.05; **, P < 0.01; ***, P < 0.001.

## Acknowledgements

This work was supported by Defense Threat Reduction Agency (DTRA) grants HDTRA1-15-1-0032 (R.B.K.) and HDTRA1-15-1-0047 (W.B.K.) and NIH grants U19 AI083019, R01 NS052632, and R01 AI101400, (R.S.K.). We have no conflicting interests to declare. Correspondence and requests for materials should be addressed to R.S.K.

---

**FIG S1.**
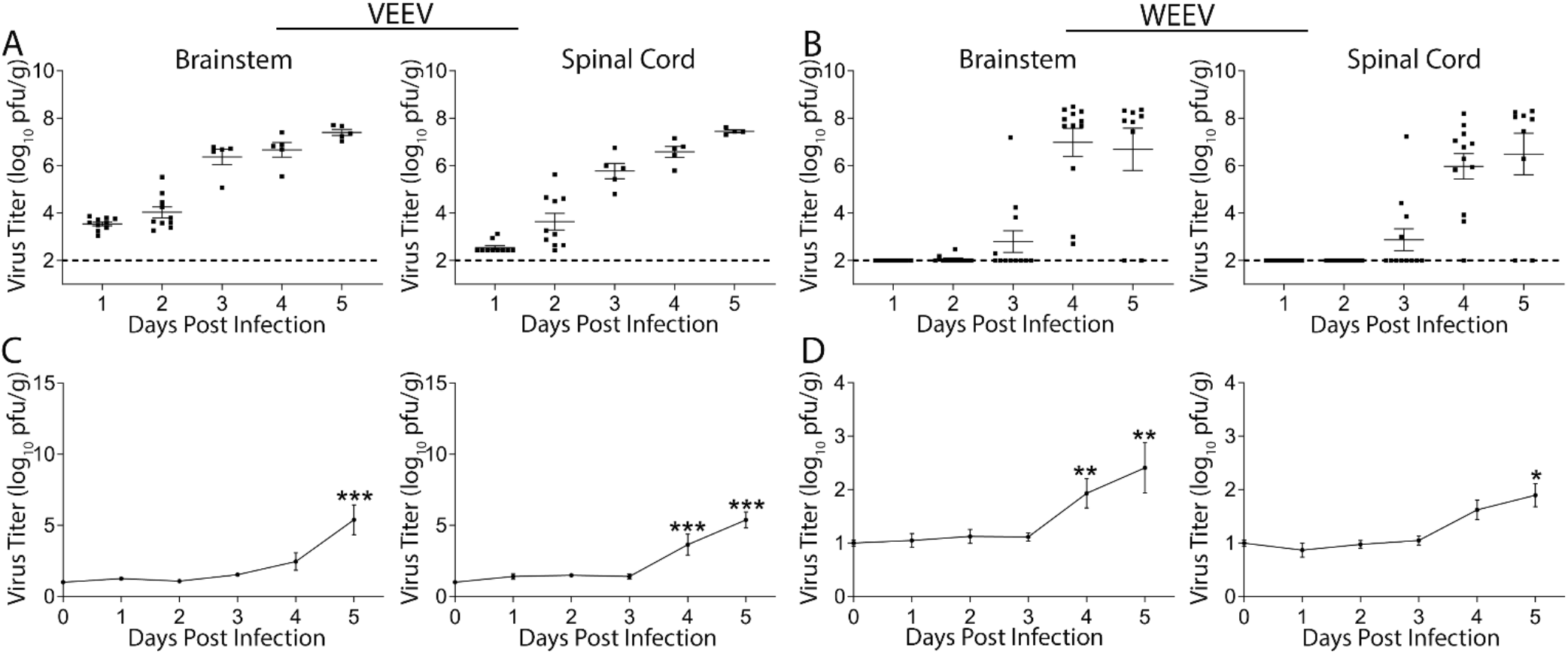
Alphavirus enters the CNS in the presence of an intact BBB. (A and B) Viral burdens in brain tissues of C57BL/6 mice following f.p. infection with either VEEV (10 PFU) or WEEV (1000 pfu) were determined via plaque assay. Dashed lines indicates detection limit of the assay. (C and D) BBB permeability was evaluated at indicated dpi by measuring sodium fluorescein in CNS tissues following i.p. injection. Results are the combined data of two independent experiments. Data presented as mean viral titer or fluorescence ± SEM for N=5-10 mice/group. *, *P* < 0.05, **; *P* < 0.01; ***, *P* < 0.001, via 1-way ANOVA.

**FIG S2.**
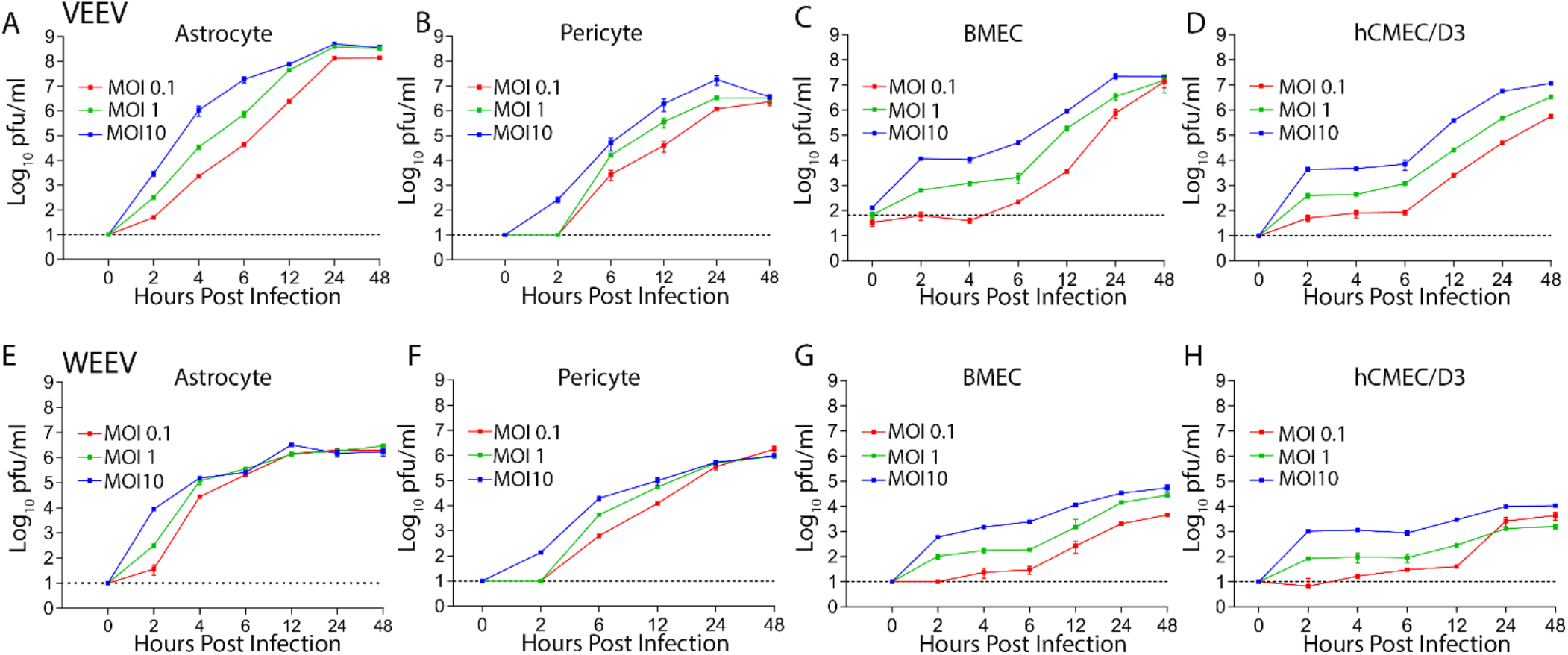
Cells of the neurovascular unit are permissive to alphavirus infection *in vitro.* Replication kinetic of VEEV and WEEV in murine (A-C, and E-G) and human cells (D and H) of the NVU. Multi-step growth curves were generated using Graphpad Prism 7. Shown is a representative data from three independent experiments each in duplicates.

**FIG S3.**
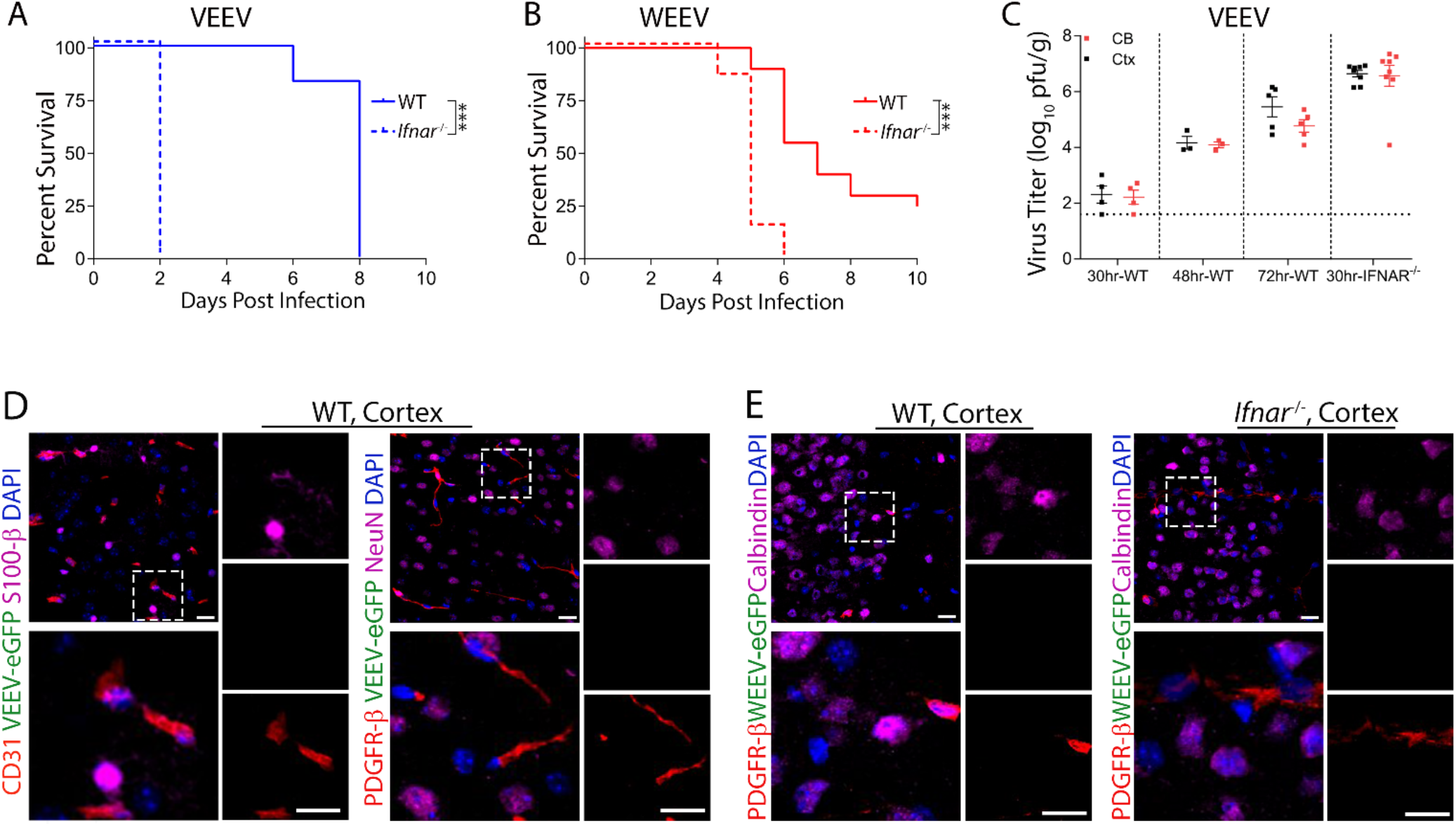
Survival phenotype of WT and *Ifnar^-/-^* mice following alphavirus infection. (A and B) WT and *Ifnar^-/-^* mice were infected with VEEV (10 pfu) and WEEV (1000 pfu) via foot-pad injection. Infected animals were monitored for survival and weight loss for 20 days. Shown is the combined data from 2 independent experiments with 3-4 animals each time. (C) Viral titers in the brain of WT versus *Ifnar^-/-^* following infection with VEEV-eGFP (100 pfu) at indicated time-point. (D and E) IHC examining of brain tissues collected from infected WT (D) and *Ifnar^-/-^* (E) mice at 1 dpi. Cell markers: astrocytes (S100-β), BMECs (CD31), pericytes (PDGFR-β) and neurons (NeuN). Nuclei counterstained with DAPI (blue). Magnification 40x. Scale bar: 20 µm.

**FIG S4.**
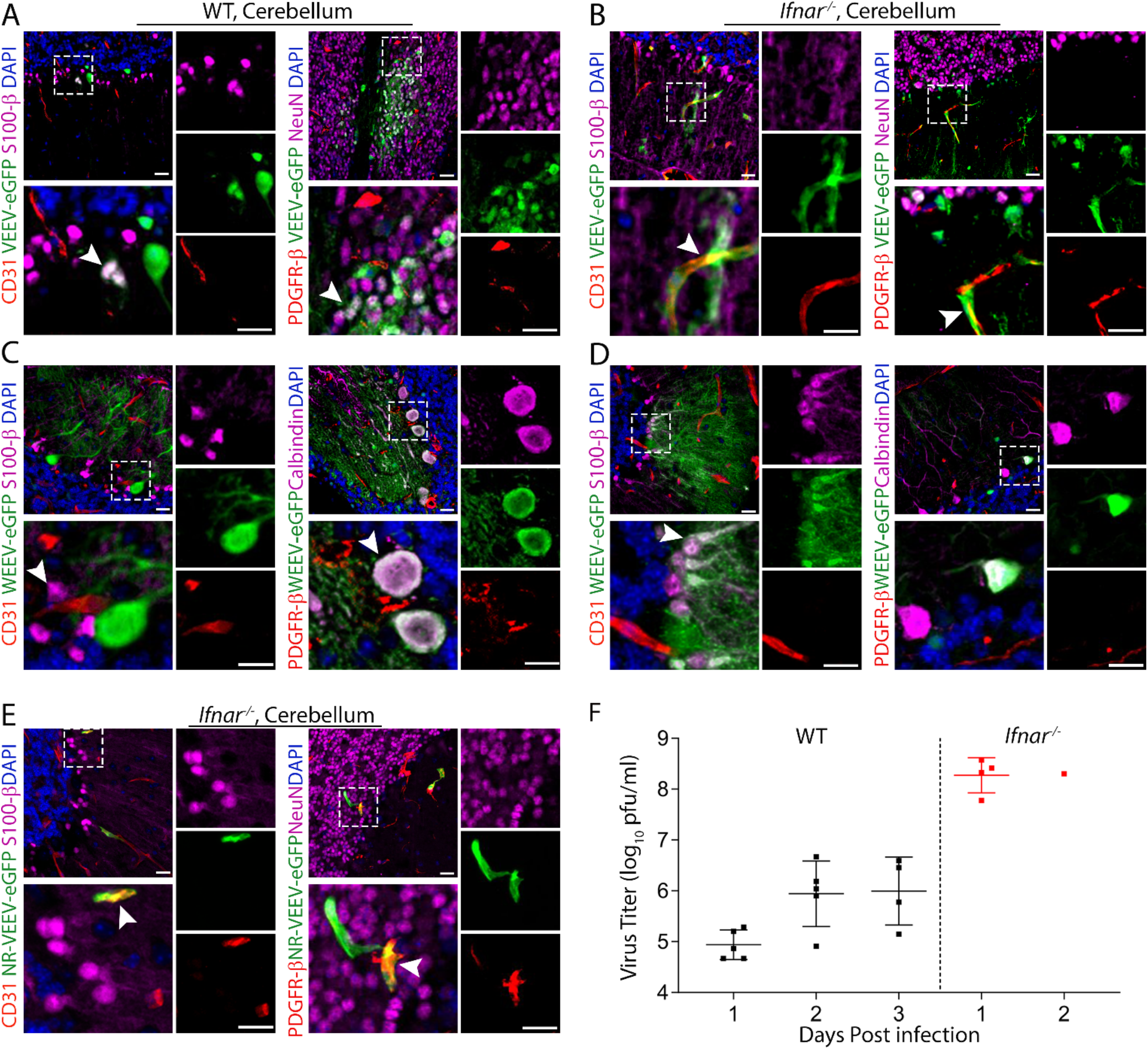
IFN-mediated restriction of alphavirus within NVU is uniform across the brain. IHC staining of brain tissues from WT (3dpi; A and C) and *Ifnar*^-/-^ (2 dpi; B and D) mice infected with either 100 pfu of VEEV-eGFP (A and B) or 1000 pfu of WEEV-eGFP via f.p. injection (C and D). Cerebellum sections were stained for markers of astrocytes (S100-β), BMECs (CD31), pericytes (PDGFR-β) and neurons (NeuN or Calmodulin). Nuclei counterstained with DAPI (blue). Magnification 40x. Scale bars: 20 µm. (E) Infection of cerebral BMECs and pericytes in *Ifnar*^-/-^ mice following *i.v.* infection with a non-replicative (NR) replicon of VEEV-eGFP. (F) Serum titers of VEEV in WT versus IFNAR^-/-^ mice following f.p. infection.

**FIG S5.**
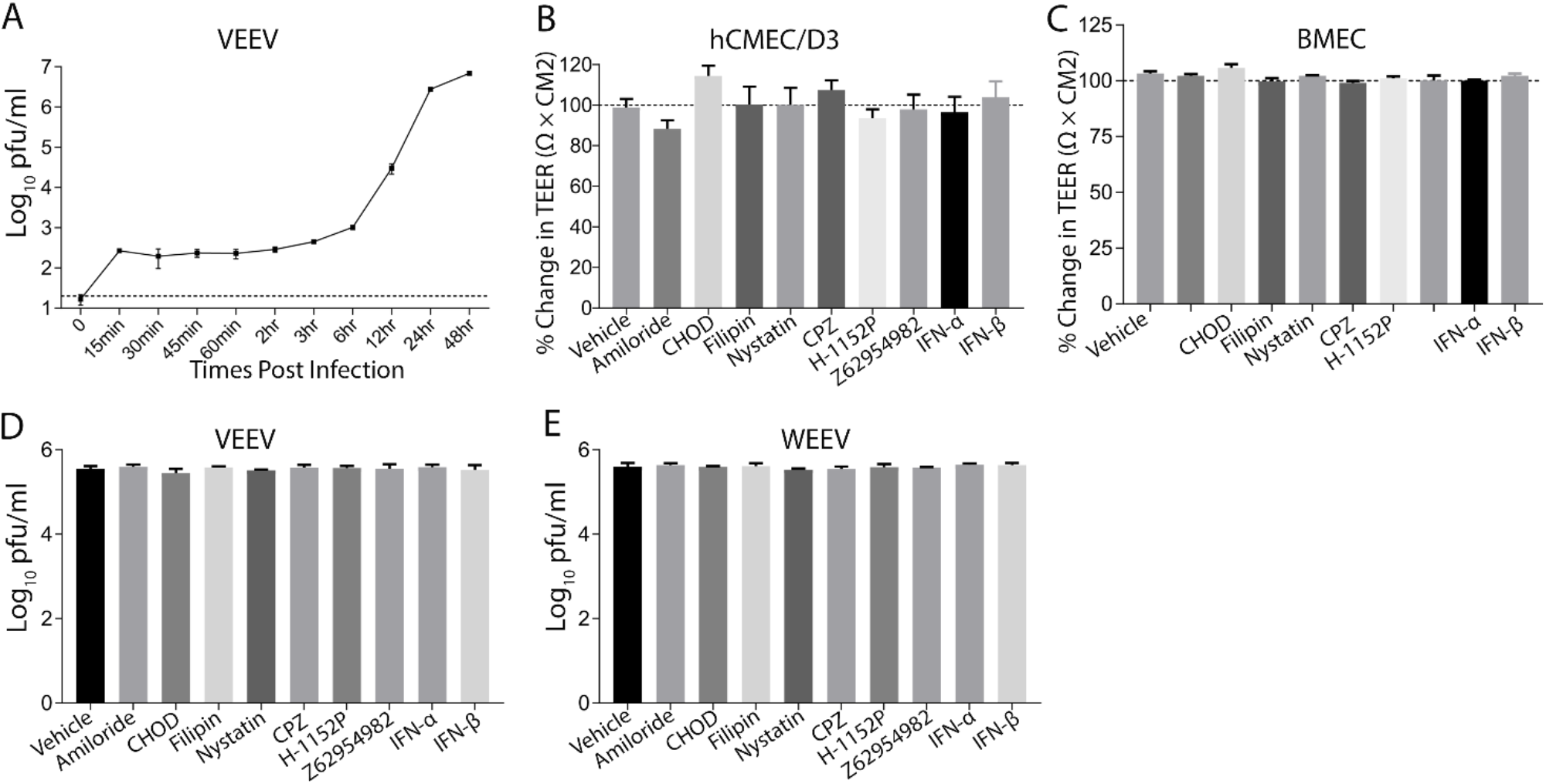
Replication kinetics of VEEV in BMECs. (A) BMECs were infected with VEEV at a MOI of 2 for 30 min. After extensive washes, virus production was assayed in the culture medium at indicated time-points using plaque assay. (B and C) Effects of inhibitors of endocytic pathways on barrier integrity, which was determined by measuring TEER before and after addition of inhibitors to BMECs on the top chamber of a transwell plate. (D and E) Effects of endocytosis inhibitors on virus infectivity. Viruses were incubated with inhibitors or vehicle for 1 hr, followed by assessment of virus infectivity using plaque assay in BHK-21 (VEEV) or Vero (WEEV) cell lines. Drug concentrations: Amiloride hydrochloride hydrate (50 µm), Cholesterol oxidase (CHOD, 2 U/ml), filipin (1 µg/ml), nystatin (12 µg/ml), chlorpromazine (CPZ, 10 µg/ml), H-1152P (50 nm, 100 nm), Z62954982 (100 µM), IFN-α (100 pg/ml), IFN-β (100 pg/ml).

**FIG S6.**
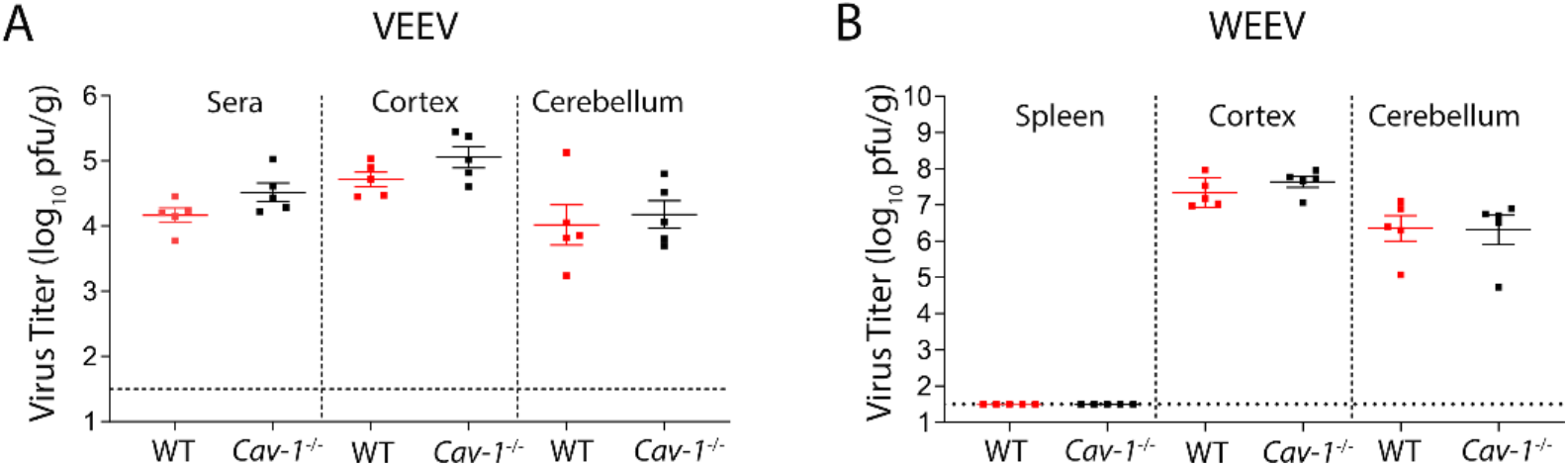
Difficinecy in *Cav-1* does not affect alphavirus replication efficiency within the CNS. (A and B) Viral titers in peripheral and brain tissues of WT vs *Cav-1*^-/-^ mice at 1dpi following i.c. infection with VEEV (10 PFU) and WEEV (100 pfu). Error bars indicate standard error of the mean (SEM).

